# The transcriptional changes underlying the flowering phenology shift in response to climate warming

**DOI:** 10.1101/2023.01.24.524815

**Authors:** Hideyuki Komoto, Yuki Hata, Junko Kyozuka, Yui Kajita, Hironori Toyama, Ai Nagahama, Akiko Satake

**Affiliations:** Department of Biology, Faculty of Science, Kyushu University, Fukuoka 819-0395, Japan; Graduate School of Life Sciences, Tohoku University, Sendai 980-8577, Japan; Iriomote Station, Tropical Biosphere Research Center, University of the Ryukyus, Okinawa, 907-1541, Japan; Biodiversity Division, National Institute for Environmental Studies, Tsukuba, Ibaraki, 305-8506, Japan; Department of Botany, National Museum of Nature and Science, 4-1-1 Amakubo, Tsukuba, Ibaraki 305-0005, Japan

**Keywords:** *Arabidopsis*, climate warming, phenological shift, molecular phenology, vernalization, *FLOWERING LOCUS C*

## Abstract

Climate warming is causing shifts in key life-history events, such as flowering time. To assess the impacts of climate warming on flowering phenology, it is crucial to understand the transcriptional changes of genes underlying the phenological shifts. Here, we comprehensively explore genes that contribute to the flowering phenology shifts under climate warming by monitoring the latitudinal gradient of seasonal expression dynamics of 293 flowering-time genes in a perennial herb, *Arabidopsis halleri*. Through transplant experiments conducted at northern, southern, and subtropical study sites in Japan, we have demonstrated that the flowering period was shortened with decreasing latitude and ultimately, the plants lost the opportunity to flower in subtropical climates. The key transcriptional changes underlying the shift in flowering phenology and the loss of flowering opportunity were the diminished expression of floral pathway integrator genes and genes in the gibberellin synthesis and aging pathways, all of which are suppressed by increased expression of *FLOWERING LOCUS C*, a central repressor of flowering. These results suggest that the upper-temperature threshold beyond which plants cannot reproduce in response to warming is governed by a relatively small number of genes that suppress reproduction in the absence of winter cold.

## 1 Introduction

The life cycles of organisms are largely driven by temperature-related cues, resulting in profound changes in biological systems under climate warming (Rosenzweig et al., 2008). The phenological shifts in key life-history events have been suggested to occur in wild plants and animals in response to warming (Collins et al., 2021; Stuble et al., 2021; Vitasse et al., 2021; Bates et al., 2022). The flowering phenology, which is a crucial life-history component and has a significant impact on reproductive success (Kudo, 1993; Munguía-Rosas et al., 2011; Anderson et al., 2012), is regulated by seasonal temperature changes. The molecular mechanisms of the response to seasonal temperature changes have been explored in laboratory-based model systems (Sung and Amasino, 2005; Bäurle and Dean, 2006; Andrés and Coupland, 2012). Vernalization is one of the most well-studied responses to seasonal temperature change at the molecular level. In *Arabidopsis*, before the winter, the floral suppressor *FLOWERING LOCUS C (FLC)* (Michaels and Amasino, 1999) down-regulates the floral activator *FLOWERING LOCUS T (FT)*, a component of the mobile flower-promoting signal (florigen) (Kardailsky et al., 1999; Kobayashi et al., 1999; Turck et al., 2008; Putterill and Varkonyi-Gasic, 2016). After exposure to the prolonged cold in winter, *FLC* is epigenetically silenced, resulting in *FT* induction and flowering in spring (Amasino, 2010; Andrés and Coupland, 2012; Whittaker and Dean, 2017).

To assess the susceptibility of plant phenology to climate change, a cross-scale approach that connects phenological shifts occurring in natural ecological systems to the underlying molecular mechanisms is promising (Satake et al., 2022). The molecular mechanism of flowering time, uncovered in laboratory-based model systems, can be applied to plants in natural conditions by monitoring seasonal changes in gene expression levels, called molecular phenology (Kudoh, 2016). This approach revealed that, even in a fluctuating natural environment, 83% of *FLC* expression variation can be explained by the temperatures over the preceding six weeks in *Arabidopsis halleri* subsp. *gemmifera* (Brassicaceae, hereafter referred to as *A. halleri)*, a perennial relative of the annual *A. thaliana* (Aikawa et al., 2010). Furthermore, they have predicted and empirically supported that the flowering period will be shortened under warming conditions as a result of an advanced shift in the reversion to vegetative growth based on the model focusing on the seasonal gene expression dynamics of *AhgFLC-AhgFT* involved in vernalization (Satake et al., 2013; Nagahama et al., 2018). While the simple model is useful for assessing vulnerability to warming, the regulation of flowering phenology shifts can be more complex and may not be fully explained by the vernalization pathway alone.

Flowering time is regulated by a complex network of multiple pathways comprising more than 300 genes (Pajoro et al., 2014; Blümel et al., 2015; Bouché et al., 2016). In addition to the vernalization pathway, genes involved in the gibberellin synthesis (Mateos et al., 2015) and aging pathways (Deng et al., 2011; Madrid et al., 2021) have been shown to be controlled by *FLC*, providing another potential mechanism for mediating the temperature-dependence of flowering phenology. However, the possibility of such mechanisms operating in natural conditions has yet to be comprehensively examined, and the genes involved in the regulation of flowering phenology shifts in response to climate warming remain largely unknown.

In this study, we comprehensively explore genes that contribute to flowering phenology shifts under climate warming by monitoring the seasonal expression dynamics of 293 flowering-time genes and flowering phenology of *A. halleri* in three transplanted sites at different latitudes: a northern site (Sendai), a southern site (Fukuoka), and a subtropical site (Iriomote Island) in Japan (Fig. 1a). By identifying genes that exhibit different seasonal expression dynamics across the three different latitudes, we demonstrate that *AhgFLC* is a central regulator of vernalization, gibberellin synthesis, and aging, influencing the flowering phenology shifts under climate warming in *A. halleri*.

**FIGURE 1.**
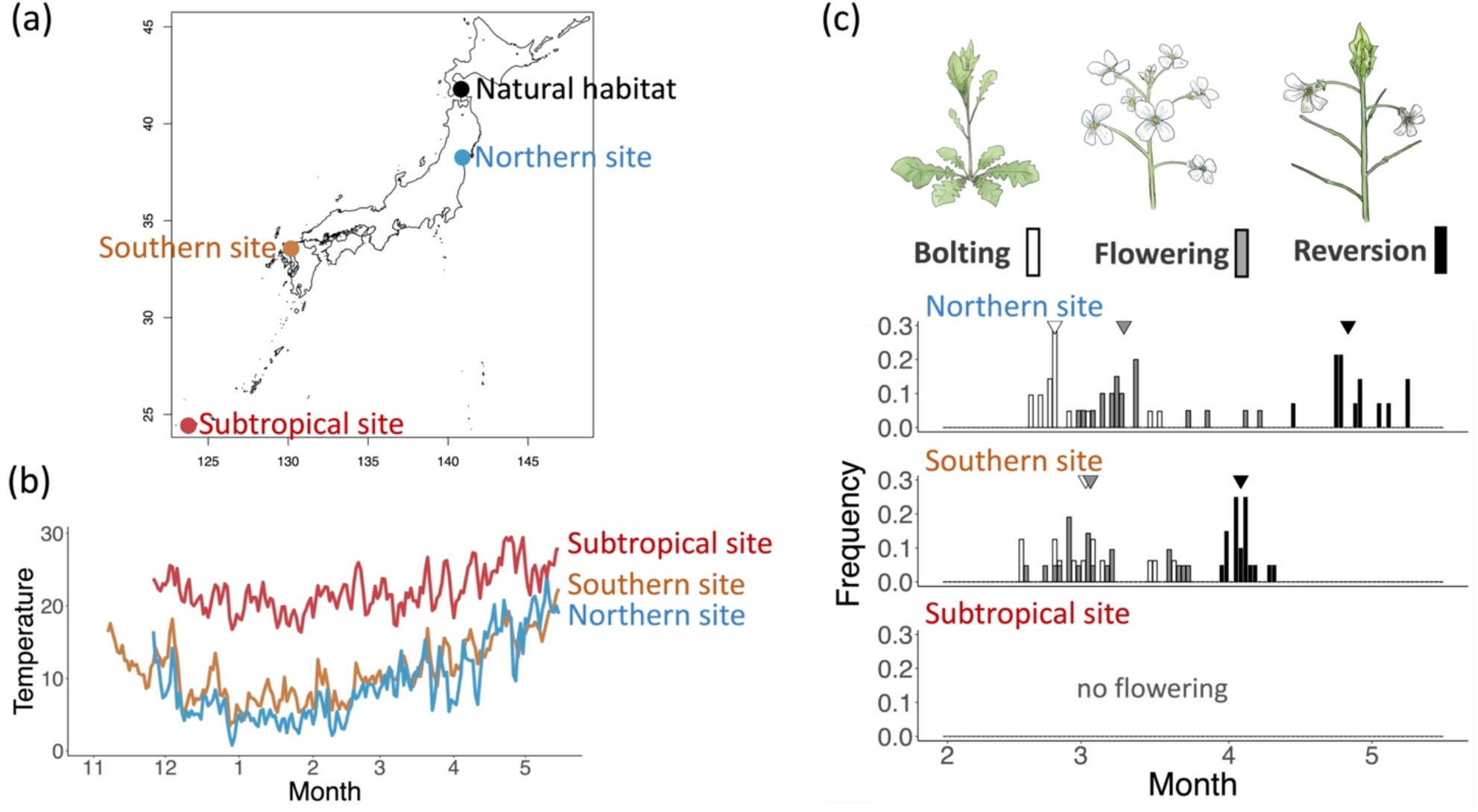
Temperature and flowering phenology of transplanted 3 sites at different latitudes. (a) The natural habitats of the Hakodate populations as well as the locations of the northern site (Sendai), the southern site (Fukuoka), and the subtropical site (Iriomote island) for the transplant experiments. (b) Dynamics of the daily average temperature at the three transplanted sites. (c) Phenology of *A. halleri* at the three transplanted sites. The inverted triangle symbols indicate the median onset dates for bolting, flowering, and reversion in each study site.

## 2 Materials and Methods

### 2.1 Natural habitat and transplant experiments

*A. halleri* (L.) O’Kane & Al-Shehbaz subsp. *gemmifera* (Matsum.) O’Kane & Al-Shehbaz is a wild perennial species closely related to *A. thaliana* and distributed in the Russian Far East and East Asia (Hoffmann, 2005). We collected 30 individuals from the natural population in Hakodate, Hokkaido, a northern island of Japan (along the Matsukura River, Hakodate City; 41°47’ N, 140°49’ E, alt. 10 m; see Fig. 1a) on 11 June 2018. The mean air temperature in Hakodate was 9.1°C (Japan Meteorological Agency 1981–2010) (Satake et al., 2013). Because it was difficult to prepare genetically uniform seeds of this self-incompatible species, we used areal rosettes formed by clonal reproduction of parent lines. The plants were grown under a continuous white fluorescent light at 22°C until clonal reproduction (formation of aerial rosettes) and aerial rosettes of about 3 cm were collected from the parent lines after flowering and reversion.

A total of 141 individuals, comprising 118 aerial rosettes (mean ± sd of rosette size = 4.79 ± 1.73 cm) from 9 parental lines and 23 plants (mean ± sd of rosette size = 2.15 ± 0.65 cm) germinated from the cross-pollinated seeds from parent lines (Suppl. Table. 1), were planted in a 7.5-cm-diameter pot containing a 6:1 (v/v) mixture of vermiculite and culture soil. These plants were grown under continuous white fluorescent light at 22°C for about 4–8 weeks before transplantation.

Among 141 individuals, we selected sufficiently grown 126 individuals, comprising 106 from aerial rosettes from 9 parent lines and 20 from cross-pollinated seeds. We transplanted these individuals to the three common gardens at different latitudes: the northern site (the common garden on Katahira campus of Tohoku University, Sendai, 38°16’N, 140°54’E, alt. 39 m; Fig. 1a), the southern site (the common garden on Ito campus of Kyushu University, Fukuoka, 33°34’N, 130°11’E, alt. 5 m; Fig. 1a), and the subtropical site (the common garden in a greenhouse depending on the ambient temperature without additional heating, on Iriomote Station, Tropical Biosphere Research Center, University of the Ryukyus, 24°26’N, 123°46’E, alt. 10 m; Fig. 1a) to monitor the molecular and flowering phenology (Suppl. Table. 1). The plants originated from different parental lines or cross-pollinated seeds were almost evenly allocated to the three common gardens: 36 (mean ± sd of rosette size = 7.00 ± 1.57 cm), 40 (6.16 ± 2.75 cm), and 50 (7.45 ± 2.06 cm) plants were used for each site (Suppl. Table. 1). The pots were placed at least 20 cm apart and were placed so that local sunlight and temperatures would be unbiased by parental lines.

Throughout the experiment, the soil surface temperature was recorded every hour at the three transplantation sites, using a HOBO H8 Pro temperature logger (Onset Computer Corporation). At all sites, the logger was placed just beneath the surface of the soil in the common gardens. Other environmental variables, such as day length, sunshine duration, and precipitation, were obtained from the Calendar Calculation—National Astronomical Observatory of Japan (https://eco.mtk.nao.ac.jp/cgi-bin/koyomi/koyomix.cgi) and Japan Meteorological Agency (https://www.data.jma.go.jp/gmd/risk/obsdl/index.php) respectively.

### 2.2 RNA-seq analysis

From November 2018 to April 2019, young leaves were sampled from three individuals every 3–5 weeks, six hours after dawn in each common garden (Fig. 2a; Suppl. Table. 2). The first sampling in November 2018 was conducted in a laboratory setting. The leaf samples were immediately preserved in a 2 ml microtube containing 1.5 ml of RNA-stabilizing reagent (RNAlater; Ambion, Austin, TX, USA). The samples were transferred to the laboratory within one hour of sampling, stored at 4°C overnight, and then stored at −80°C until RNA extraction. RNA integrity was examined using the Agilent RNA 6000 Nano kit on a 2100 Bioanalyzer (Agilent Technologies, Santa Clara, CA, USA), and the RNA yield was determined using a NanoDrop ND-2000 spectrophotometer (Thermo Fisher Scientific, Waltham, MA, USA). Five to six micrograms of RNA extracted from the leaves of each sample were sent to Hokkaido System Science Co., Ltd., where a cDNA library was prepared with a NovaSeq 6000 S4 Reagent Kit, and paired-end transcriptome sequencing of each sample was conducted using an Illumina NovaSeq6000 sequencer (Illumina, San Diego, CA, USA). Over a total of 30 million 101 base paired-end reads were obtained (Suppl. Table. 2).

**FIGURE 2.**
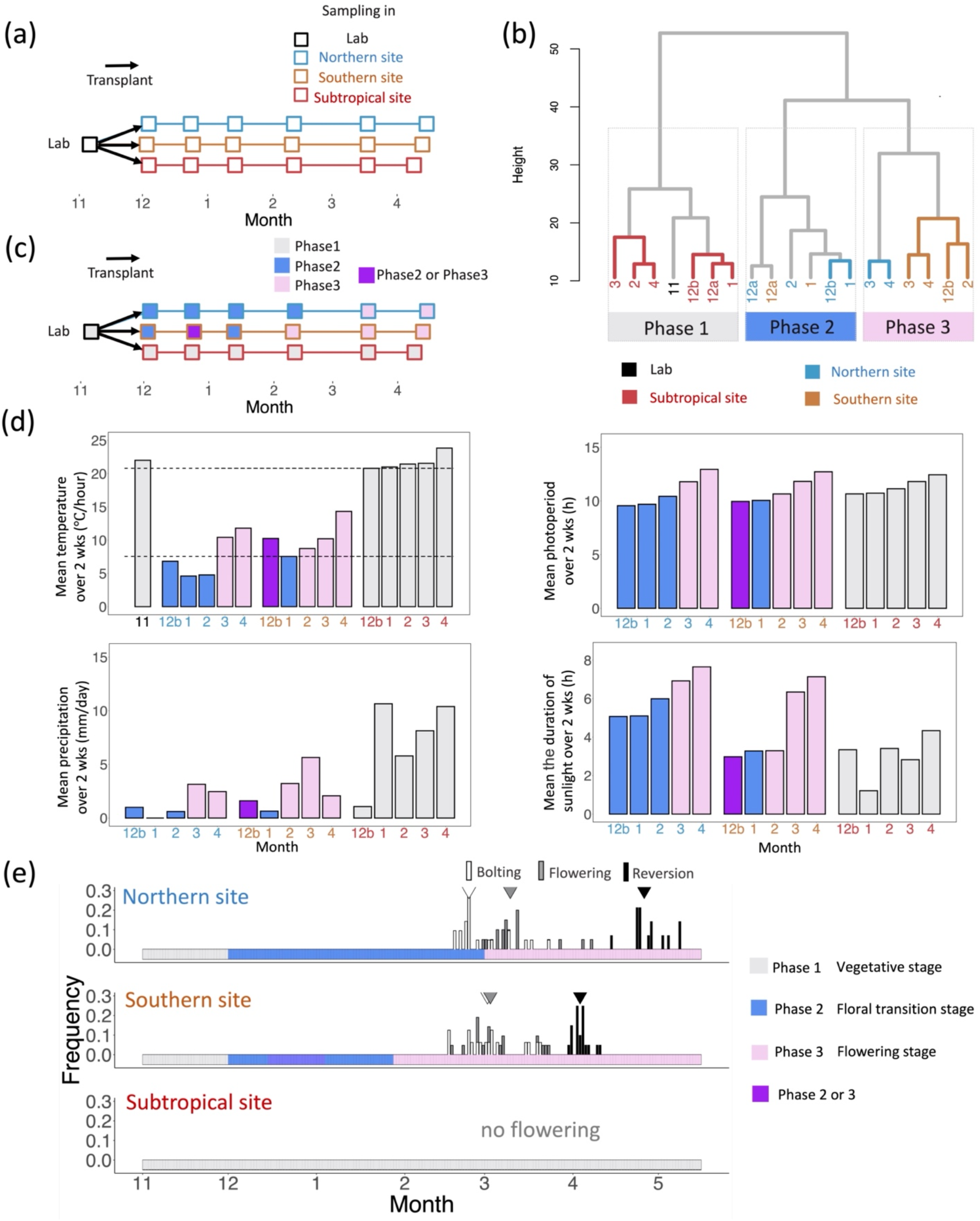
Association between molecular phenology and seasonal changes of environmental variables and the stage of floral development. (a) Sampling schedule at the lab and three transplanted sites. The plants were transplanted from the laboratory to three different latitude sites in early December and sampled every 3–5 weeks at each site. (b) Hierarchical clustering of monthly transcriptome profiles. The number indicates the month when the RNA was extracted. When sampling was performed twice a month, the month is distinguished by the letters “a” or “b”. Gene expression profiles were divided into three phases: phase 1 (gray), phase 2 (blue), and phase 3 (pink). (c) The phase of each sample is based on hierarchical and non-hierarchical clustering. The phase shown in purple indicates that the results are different in the two clustering methods. (d) The mean temperature, photoperiod, precipitation, and duration of sunlight over two weeks before the sampling dates. Dashed lines indicate 20.7°C, above which temperature the phase 1 profiles in gene expression emerge, and 6.76°C, below which temperature the phase 2 profiles in gene expression emerge. Note that the plants were not directly impacted by rainfall as they were shielded by a roof at the subtropical site. (e) Correspondences of molecular phenology of phase 1(gray), phase 2 (blue), phase 2 or 3 (purple), and phase 3 (pink) to the stage of floral development. The inverted triangle symbols indicate the median onset dates for bolting, flowering, and reversion in each study site.

The quantification of expression was conducted as follows. First, quality filtering was performed using the fastp v0.20.0 with default options (Suppl. Table. 3). Second, RNA-seq reads were mapped to the reference genome (Briskine et al., 2017) using STAR 2.7.3a (Suppl. Table. 4). Third, RNA-seq reads were quantified using RSEM v1.3.1.

### 2.3 Records of flowering phenology

The dates of onset of bolting, flowering, and reversion were recorded following a previous study (Satake et al., 2013). The onset of bolting was defined as the elongation of floral stems beyond 2.0 cm. The onset of reversion was defined as the emergence of aerial rosettes with a diameter greater than 0.5 cm on apical or lateral meristems, or senescence of the apical meristem (Nagahama et al., 2018). Aerial rosettes were not always produced, and basal rosettes produced new leaves after flowering. To ensure a sufficient sample size for monitoring flowering phenology, phenological observations were conducted on all transplanted plants, regardless of whether they were used in gene expression analysis. The median onset dates for bolting, flowering, and reversion in each study site were calculated with 95% confidence intervals using 1,000 bootstrap resamples.

### 2.4 Seasonality in transcriptional profiles of flowering-time genes

We selected 23,400 genes with TPM (Transcripts Per Million) more than 1 in at least one sample as expressed genes. Among 23,400 expressed genes, we further extracted genes associated with flowering time regulation using a database as reported in *A. thaliana* (Bouché et al., 2016) to capture the molecular phenology that primarily contributes to flowering phenology. The database comprises information about 306 genes, published in 1,646 articles authored by 4,606 scientists. We searched for homologs of each *A. halleri* gene using BLAST searches (blastp) for all coding sequences in *A. thaliana*. We used the genes with the best hit and with an e-value less than 1e-5. Among the 306 flowering-time genes in *A. thaliana*, 293 were identified as expressed genes in *A. halleri*.

For the statistical analyses, TPM of those genes were transformed to log2[TPM+0.01] to normalize variations in gene expression and the mean across three biological replicates was calculated for each time point in each study site. To assess the seasonality in the transcriptional profiles of these genes, we first performed hierarchical and non-hierarchical clustering of 19 transcriptional profiles, using the Ward method and the *K*-means method, respectively, based on the Euclidian distance in gene expression level. These clustering methods were implemented using the hclust and kmeans functions in R (ver. 3.6.3). To determine the number of clusters in non-hierarchical clustering, the elbow method was applied (Suppl. Fig. 1a).

We then conducted a principal component analysis (PCA) of the gene expression level from all samples using the prcomp function of the stats package in R (ver 3.6.3). To identify the genes and characteristics that primarily contribute to the first principal component (PC1) and the second principal component (PC2), we extracted the top 5 % genes with large absolute eigenvectors, in PC1 (Supplementary Table 5) and PC2 (Supplementary Table 6), respectively. The axis of the third principal component (PC3), which distinguished between the sample of March and April in the northern site and the others (Suppl. Fig. 2), was not considered in this analysis as its relationship to flowering phenology was ambiguous.

### 2.5 Visualizing the seasonal transcriptional changes in the network of flowering-time genes

To identify the key genes and regulatory pathways associated with the flowering phenology shifts, we first draw a flowering gene network and then analyzed the shifts in major transcriptional changes across three phases determined by two clustering methods. The three phases are associated with the response to seasonal temperature change and the stages of floral development. We draw the flowering gene network using the database based on the knowledge of *A. thaliana* (Bouché et al., 2016) using the top 5 % genes with large absolute eigenvectors in PC1. The regulatory relationship between *FLC* and genes related to hormonal pathways was also based on another study (Mateos et al., 2015) (Suppl. Table. 7). Four genes with no regulatory relationship among the extracted genes were omitted from the network (Suppl. Table. 5 and 7). To identify the pairwise regulatory relationships among genes comprising the flowering gene network based on expression levels and to clarify whether these relationships are consistent across *A. halleri* and *A. thaliana*, we performed linear regression analysis and calculated the slope with p-value and 95% confidence interval using the lm function (Suppl. Table. 7). Additionally, we identified genes that were differentially expressed in phase 2 and 3 compared to phase1 among genes comprising the flowering gene network by conducting a student t-test (two-sided) with Bonferroni correction using the t.test function in R (ver. 3.6.3).

## 3 Results

### 3.1 The latitudinal gradient of flowering phenology and the loss of flowering opportunity in the subtropical site

The temperature from November 2018 to April 2019 in the southern and the subtropical site was 1.99°C and 13.68°C higher than in the northern site, respectively (Fig. 1b). Our transplant experiments showed that flowering period became shorter as latitude declined (Fig. 1c), and the plants completely lost the flowering opportunity in the subtropical site. The flowering period in the southern site, measured as the period from the median date of bolting to that of reversion, was 29 days shorter than that in the northern site (Fig. 1c). The median (95% confidence interval) of bolting, flowering, and reversion dates in the northern site were 23 February (22 February–2 March), 10 March (8 March–13 March), 26 Apr. (24 April–1 May), respectively. In contrast, those in the southern sites were 1 March (24 February–9 March), 2 March (26 February–7 March), and 3 April (2 April–4 April), respectively. In the subtropical site, we continued monitoring for one more year and confirmed that no flowering occurred for two consecutive years.

### 3.2 Molecular phenology of flowering-time genes was associated with seasonal changes in temperature and the stage of floral development

Hierarchical clustering of our field transcriptome data from the three study sites revealed that transcriptional profiles were classified into three phases (Fig. 2b). Non-hierarchical clustering showed similar results, except that the sample collected in late December at the southern site belonged to phase 2 (Fig. 2bc; Suppl. Fig. 1b).

The transcriptional profile of phase 2 was observed when the mean temperature over the 2 weeks preceding the monitoring date was below 6.76°C (Fig. 2d), while phase 1 emerged when the mean temperature exceeded 20.7°C. Between these temperature thresholds, phase 3 was observed (Fig. 2d). Irrespective of the window size employed to calculate the mean temperature, the distinction among the three phases was associated with variations in temperature (Suppl. Fig. 3). Other environmental variables such as the photoperiod, duration of sunlight, and precipitation, were not strongly correlated with seasonal transcriptional profiles (Fig. 2d).

The three phases classified by the clustering methods were not only associated with the response to seasonal temperature changes but also the stages of floral development. Transcriptional profiles of phase 1 appeared during a vegetative stage (Fig. 2e). Transcriptional profiles shifted from phase 1 to phase 2 in December at the northern and southern study sites and bolting occurred at the end of or after one month later of phase 2 in these study sites (Fig. 2e). Therefore, the transcriptional profile of phase 2 is likely to characterize the floral transition stage. As the season progressed, transcriptional profiles transitioned from phase 2 to phase 3 in March at the northern site and at the end of January at the southern site (Fig. 2e). Because flowering was observed during phase 3, the transcriptional profile of phase 3 corresponds to the flowering stage (Fig. 2e).

### 3.3 The loss of flowering opportunity in the warmest environment can be explained by silenced *AhgFT* and *AhgTSF*

PCA showed that the first two axes of variation (the principal components; PCs) accounted for 56.3% of the variation in the multidimensional functional space (Fig. 3ab). The axis of the PC1 captured the differences in gene expression profiles between the subtropical site and the other sites (Fig. 3ab). In contrast, the axis of the PC2 distinguished gene expression profiles between the northern site and the others (Fig. 3ab).

**FIGURE 3.**
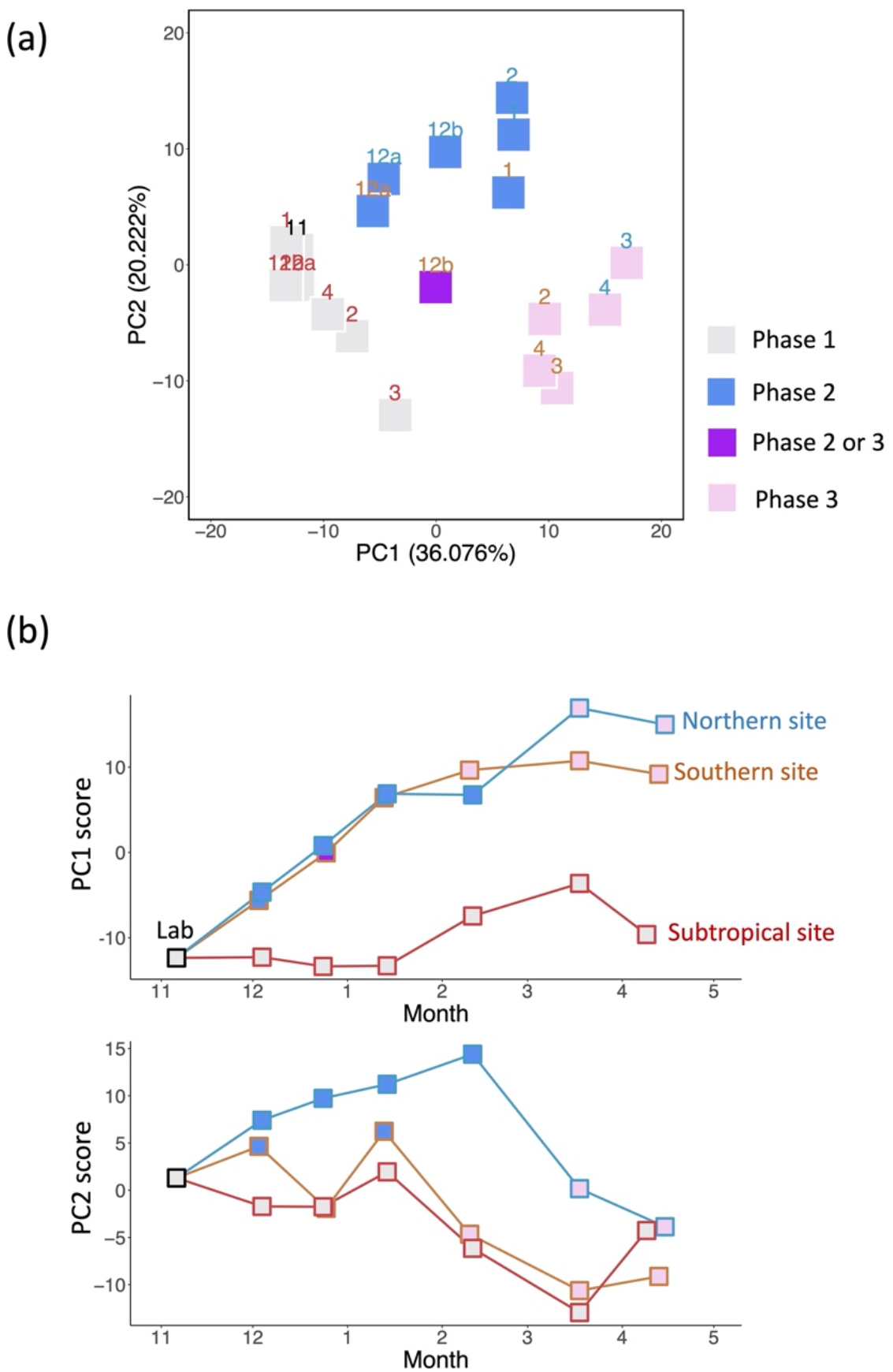
Major axes of multivariate molecular phenology. (a) A plot of PC2 versus PC1 resulting from the PCA of 293 genes. Letter colors indicate sampling sites: the lab (black), the northern site (blue), the southern site (orange), and the subtropical site (red). Plot colors indicate the gene expression profiles for phase 1(gray), phase 2 (blue), phase 2 or 3 (purple), and phase 3 (pink). The numbers in brackets represent the explained variance. (b) Plots of the PC1 and PC2 scores against the month.

The top 5% of genes with large absolute eigenvectors of each PC include 15 genes (Suppl. Table 5 and 6). Among the top 5% genes for both PCs, *AhgFLC, AhgSPL15 (SQUAMOSA PROMOTER BINDING PROTEIN-LIKE 15), AhgTEM2 (TEMPRANILLO 2)*, and *GA2ox6* (*GIBBERELLIN 2-OXIDASE 6*) were overlapped (Suppl. Table 5 and 6).

*AhgFT* and *AhgTSF* (*TWIN SISTER OF FT)* exhibited the highest absolute eigenvectors of PC1 (Fig. 4a). The expression levels of these genes were low throughout the census in the subtropical site, while those in the other sites were elevated from winter to spring (Fig. 4b). Similar expression profiles were observed in genes included in the top 5% genes with positive eigenvectors of PC1 (Fig. 4c). The expression levels of genes with the negative eigenvectors exhibited an opposite trend (Fig. 4c). The distinct expression profiles in the subtropical site compared to those in the other sites can explain the absence of flowering only in the subtropical site (Fig. 1c).

**FIGURE 4.**
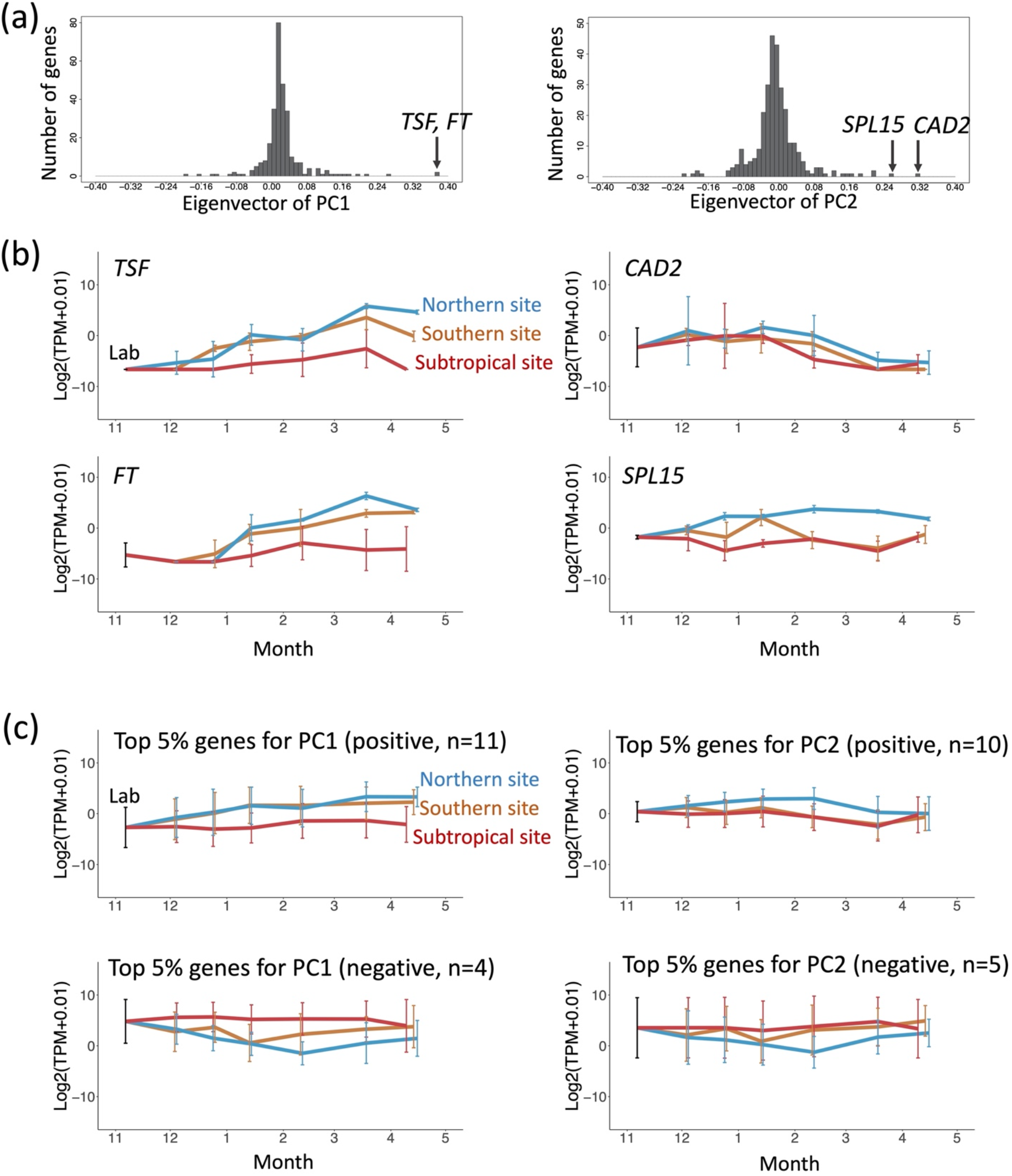
Seasonal expression profiles of genes characterizing each of the two axes. (a) Histogram of the values of eigenvectors of PC1 and PC2. (b) The expression profile of two representative genes. The two genes were selected for each of the two axes as examples: PC1 (*AhgTSF* and *AhgFT*) and PC2 (*AhgCAD2* and *AhgSPL15*). The average (line) ± s.d. (error bar) of each gene (*n* = 3; biological replicates) is shown. (c) The expression profile of the genes, specifically of those genes with either positive or negative eigenvectors in the top 5% of the largest absolute values of eigenvectors in PC1 (suppl. Table 5) and PC2 (suppl. Table 6), respectively. The average (line) ± s.d. (error bar) of those genes is shown.

*AhgCAD2 (CADMIUM SENSITIVE 2)* and *AhgSPL15* are among the top 2 genes that exhibit the largest eigenvectors of PC2 (Fig. 4a). The expression level of *AhgSPL15* was continuously higher in the northern site than the other sites throughout the census (Fig. 4b). Although the expression dynamics of *AhgCAD2* were obscured by its large variance, the top 5% of genes with positive or negative eigenvectors of PC2 generally displayed comparable or inverse trends. (Fig. 4c). These results suggest that genes that characterize PC2 are associated with the cold stress response in the northern site.

### 3.4 *AhgFLC* is a key regulator of flowering phenology shift

In the flowering-related gene network, based on knowledge of *A. thaliana* (Mateos et al., 2015; Bouché et al., 2016), 11 of the 15 genes that characterized PC1 formed 18 pairs of relationships (Fig. 5a; Suppl. Fig. 4b; Suppl. Table. 5 and 7). Among 18 pair-wise relationship of the 11 genes, the type of regulatory relationship (activation or repression) of 16 pairs was consistent between *A. halleri* and *A. thaliana* except for *AhgTEM2-AhgGA3OX1(GIBBERELLIN 3-OXIDASE 1) and AhgFLC-AhgGA2ox6* (Suppl. Table. 7). The regulatory relationship between *AhgTEM2* and *AhgGA3OX1* was unclear (Suppl. Table. 7), and the relationship between *AhgFLC* and *AhgGA2ox6* was opposite to that in *A.thaliana* (Suppl. Table. 7). Thus, the regulatory relationships of the 11 genes were largely conserved between the annual and perennial *Arabidopsis*. The flowering gene network in *A. halleri* suggests that *AhgFLC* regulates other genes involved in the hormonal or aging pathways (Fig. 5a; Suppl. Fig. 4b).

**FIGURE 5.**
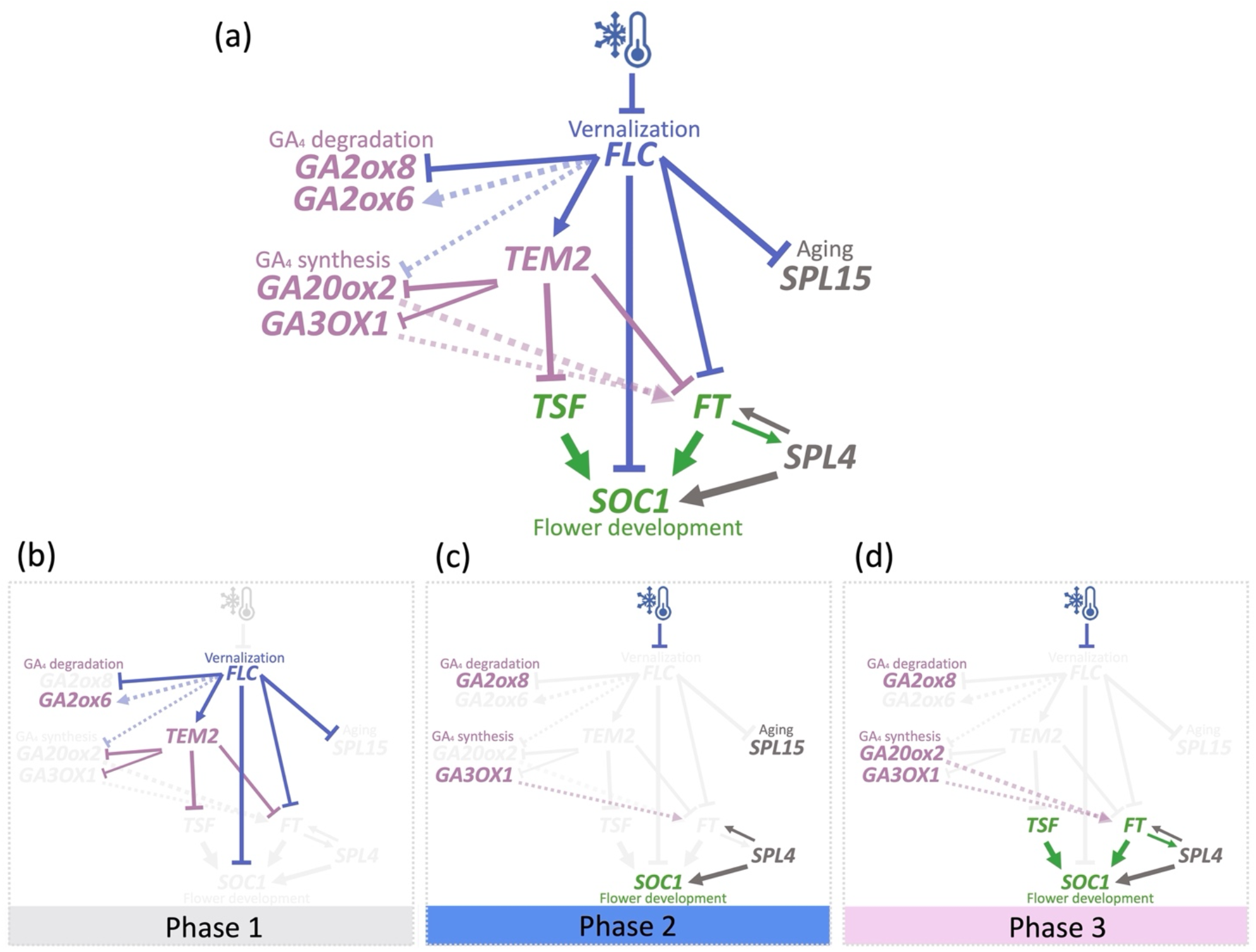
Seasonal transcriptional changes in the network of flowering-time genes. (a) Flowering gene network using the top 5 % of genes with the largest absolute values of eigenvectors for PC1. The color depends on which pathway the gene belongs to blue for the vernalization, pink for the hormonal pathway, grey for the aging pathway, and green for the flower development pathway. The dotted lines with relatively lighter colors indicate indirect effects based on the findings of *A. thaliana* (Mateos et al., 2015; Bouché et al., 2016). Arrow and T-bar indicate positive and negative regulation based on the slope of linear regression analysis. (b)–(d) Seasonal changes in gene expression comprise the flowering gene network. The significantly higher expression in each phase was shown in colored letters and symbols: in phase1 (the gene of highly expressed in phase 1 and p-value < 0.05 both between phases 1 and 2 and between phases 1 and 3) (b), in phase 2 (the gene of highly expressed in phase 2 and p-value < 0.05 between phases 1 and 2) (c) and in phase 3 (the gene of highly expressed in phase 3 and p-value < 0.05 between phases 1 and 3) (d).

To illustrate the shift of major transcriptional changes across three transcriptional and developmental phases, differentially expressed genes across three phases were highlighted in a flowering gene network (Fig. 5b–d). At the vegetative stage (phase 1), *AhgFLC, AhgTEM2*, and *AhgGA2ox6* were expressed at higher levels than in phases 2 and 3(Fig. 5b). At the floral transition stage (phase 2), the expression of *AhgGA2ox8 (GIBBERELLIN 2-OXIDASE 8), AhgGA3OX1*, *AhgSOC1 (SUPPRESSOR OF OVEREXPRESSION OF CO), AhgSPL4 (SQUAMOSA PROMOTER BINDING PROTEIN-LIKE 4)*, and *AhgSPL15* were elevated (Fig. 5c). At the flowering stage (phase 3), *AhgFT, AhgTSF*, and *AhgGA20ox2* were activated (Fig. 5d). At the northern and southern study sites, winter cold could drive the transcriptional shifts between phases 1, 2 and 3 as the season progressed. However, at the subtropical site, the transcriptional profile remained unchanged from phase 1, where floral suppressors *AhgFLC* and *AhgTEM2* were highly expressed because the warm temperature in the subtropical site was insufficient for the suppression of *AhgFLC*. These results suggest that the shift in flowering phenology is governed by the shift in transcriptional profiles and that *AhgFLC* is a key regulator for the transcriptional shift.

## 4 Discussion

We uncovered the key transcriptional changes underlying the flowering phenology shifts in a perennial *Arabidopsis*. The key transcriptional changes include the diminished expression of floral pathway integrator genes and genes in the gibberellin synthesis and aging pathways, all of which are suppressed by increased expression of *AhgFLC*. These findings contribute to deepen our knowledge regarding the molecular mechanisms underlying phenological changes in plants in response to climate change.

Our transplant experiments revealed that the flowering period was shortened as latitude decreased, and flowering opportunity was completely lost in the subtropical climate where the temperature rarely dropped below 15°C (Fig. 1bc). This is consistent with the prediction of the mathematical model, which accounts for the dynamics of *AhgFLC* and *AhgFT* expression in response to warming (Satake et al., 2013). The transcriptional changes that govern the shift in flowering phenology were best characterized by *AhgFT* and *AhgTSF* (Fig. 4a). *FT* and *TSF* are highly conserved genes that play a critical role in floral transition in angiosperm (Kardailsky et al., 1999; Kobayashi et al., 1999; Yamaguchi et al., 2005; Kinoshita and Richter, 2021). The loss of flowering opportunity is likely driven by the suppression of downstream genes, *AhgFT* and *AhgTSF*, in the gene regulatory network due to the continuous activation of *AhgFLC* during seasonal environmental changes from fall to spring (Fig. 2e; Fig. 5b; Suppl. Fig. 4b). These results suggest that the upper-temperature threshold beyond which plants are unable to adapt to global warming is controlled by a relatively small number of genes that inhibit reproduction in the absence of winter cold.

Other genes involved in gibberellin synthesis were identified as novel candidates that control flowering phenology. Gibberellins promote flowering under both short (Wilson et al., 1992) and long days (Griffiths et al., 2006; Willige et al., 2007; Galvão et al., 2012; Porri et al., 2012), and the genes that regulate gibberellins metabolism are regulated by SVP (SHORT VEGETATIVE PHASE) (Andrés et al., 2014) or SVP-FLC complex (Mateos et al., 2015) in *A. thaliana*. Given that *AhgSVP* did not exhibit clear seasonal changes in expression (Suppl. Fig. 4a) contrary to *AhgFLC* and gibberellins synthesis genes, we posit that *AhgFLC*, rather than *AhgSVP*, would be the temperature-dependent regulator of gibberellin synthesis in *A. halleri*. In phase 3, the expression of *AhgGA20ox2*, encoding the rate-limiting enzyme for gibberellin biosynthesis (Plackett et al., 2012), significantly increased compared to phase 1 (Fig. 5d). The elevated expression of *AhgGA20ox2* might contribute to the significant expression increase in *AhgFT* and onset of flowering in phase 3.

Our study also suggests that genes involved in aging are involved in the regulation of flowering phenology shift. The timing of floral transition as plant age is mediated through microRNAs and SPLs (Quiroz et al., 2021). As the plants age, the abundance of miR156 decreases, resulting in the accumulation of SPLs and floral transition (Wu et al., 2009; Hyun et al., 2016). A previous study suggested that the miR156/SPL pathway plays a role in rescuing flowering under warm conditions (Nagahama et al., 2018). However, this study did not observe such a rescue, which could potentially be attributed to the warmer environment in the subtropical site. Instead, our study revealed that expressions of *AhgSPL4* and *AhgSPL15* change in a temperature-dependent manner (Fig. 5b-d), even though the plant ages were aligned (Suppl. Table. 1). Among these SPL genes, *SPL15* is negatively regulated by *FLC* (Deng et al., 2011; Madrid et al., 2021) and flowering in response to winter cold depends on *SPL15* in perennial *Arabis alpina* (Hyun et al., 2019). Consistent with these findings, our study showed that both *AhgSPL15* and *AhgFLC* have large eigenvector values for PC2 as well as PC1 (Suppl. Table. 5 and 6), which suggests that *AhgSPL15*, as well as *AhgFLC*, plays an important role in the cold response. To fully understand the temperature dependence of *AhgSPL15* and its role in flowering time, a comprehensive analysis that includes both leaf and shoot meristem RNA samples will be required, as *SPL15* is primarily expressed in shoot meristems.

Plants have evolved diverse life histories and phenotypic plasticity in environmental responses as a result of their adaptation to the environment. The environments surrounding plants have changed continuously in the past and are changing even more rapidly under the influence of climate change in the future. Whether plants will be able to adapt to global warming in the future depends on the extent to which they can render appropriate phenotypic responses to the new environment. At least for perennial *Arabidopsis*, there exists an upper-temperature threshold beyond which the plants are unable to reproduce, resulting in an increased risk of extinction. Our approach can be applied to other species to estimate the adaptation limit and uncover central genes regulating the phenology shift. In genus Prunus, the central regulator of temperature limit appears to be the dormancy-associated MADS-box (DAM) genes that are responsible for the release of bud dormancy after winter cold (Yamane et al., 2008; Sasaki et al., 2011; Cook et al., 2012; Habu et al., 2012). Like *FLC*, most DAM genes are highly expressed before winter and downregulated in response to cold weather (Sasaki et al., 2011; Vimont et al., 2019). In the future, more data on molecular phenology will allow for a more accurate estimation of adaptation limits in diverse organisms and will advance a comprehensive understanding of the molecular mechanisms of phenological shifts in plants in response to climate change.

## 5 Acknowledgements

This work was done in support of JSPS KAKENHI Grant Number JP21H04781 to A.S. Tropical Biosphere Research Center (Iriomote Station) provided a section of the glass greenhouse for the plant cultivation. We thank K.Ota for her kind efforts in taking care of plants in the laboratory. We thank S. Kudo for his drawing of *A. halleri* in Fig. 1 (c). We thank M.shimizu for his technical support in RNA-seq analysis. We thank E. Sasaki for her helpful comments.

## 6 Author contributions

AS conceived and designed the analysis. HK has done all data analysis and interpretation. AS revised the analysis and interpretation critically. AN and AS conducted the cultivation, growth, and transplantation of the experimental plants. YH and JK conducted plant cultivation, sampling for the RNA-seq analysis, and monitoring of plant phenology at the northern site. YK and HT conducted the same tasks at the subtropical site. AN and AS conducted the same tasks at the southern site. HK wrote the manuscript, AS revised it mainly and the other authors reviewed it and provided critical feedback.

## 7 Data availability

The sequence data that support the findings of this study are available in the DDBJ Short Read Archive repository, with the accession numbers DRR437060-DRR437116.

## References

Aikawa, S., Kobayashi, M. J., Satake, A., Shimizu, K. K., and Kudoh, H. (2010). Robust control of the seasonal expression of the *Arabidopsis FLC* gene in a fluctuating environment. Proc. Natl. Acad. Sci. U. S. A. 107, 11632–11637. doi: 10.1073/pnas.0914293107.

Amasino, R. (2010). Seasonal and developmental timing of flowering. Plant J. 61, 1001–1013. doi: 10.1111/j.1365-313X.2010.04148.x.

Anderson, J. T., Inouye, D. W., McKinney, A. M., Colautti, R. I., and Mitchell-Olds, T. (2012). Phenotypic plasticity and adaptive evolution contribute to advancing flowering phenology in response to climate change. Proc. R. Soc. B Biol. Sci. 279, 3843–3852. doi: 10.1098/rspb.2012.1051.

Andrés, F., and Coupland, G. (2012). The genetic basis of flowering responses to seasonal cues. Nat. Rev. Genet. 13, 627–639. doi: 10.1038/nrg3291.

Andrés, F., Porri, A., Torti, S., Mateos, J., Romera-Branchat, M., García-Martínez, J. L., et al. (2014). SHORT VEGETATIVE PHASE reduces gibberellin biosynthesis at the *Arabidopsis* shoot apex to regulate the floral transition. Proc. Natl. Acad. Sci. 111. doi: 10.1073/pnas.1409567111.

Bates, J. M., Fidino, M., Nowak-Boyd, L., Strausberger, B. M., Schmidt, K. A., and Whelan, C. J. (2022). Climate change affects bird nesting phenology: Comparing contemporary field and historical museum nesting records. J. Anim. Ecol., 1–10. doi: 10.1111/1365-2656.13683.

Bäurle, I., and Dean, C. (2006). The Timing of Developmental Transitions in Plants. Cell 125, 655–664. doi: 10.1016/j.cell.2006.05.005.

Blümel, M., Dally, N., and Jung, C. (2015). Flowering time regulation in crops-what did we learn from Arabidopsis? Curr. Opin. Biotechnol. 32, 121–129. doi: 10.1016/j.copbio.2014.11.023.

Bouché, F., Lobet, G., Tocquin, P., and Périlleux, C. (2016). FLOR-ID: An interactive database of flowering-time gene networks in Arabidopsis thaliana. Nucleic Acids Res. 44, D1167–D1171. doi: 10.1093/nar/gkv1054.

Briskine, R. V., Paape, T., Shimizu-Inatsugi, R., Nishiyama, T., Akama, S., Sese, J., et al. (2017). Genome assembly and annotation of *Arabidopsis halleri*, a model for heavy metal hyperaccumulation and evolutionary ecology. Mol. Ecol. Resour. 17, 1025–1036. doi: 10.1111/1755-0998.12604.

Collins, C. G., Elmendorf, S. C., Hollister, R. D., Henry, G. H. R., Clark, K., Bjorkman, A. D., et al. (2021). Experimental warming differentially affects vegetative and reproductive phenology of tundra plants. Nat. Commun. 12, 1–12. doi: 10.1038/s41467-021-23841-2.

Cook, B. I., Wolkovich, E. M., and Parmesan, C. (2012). Divergent responses to spring and winter warming drive community level flowering trends. Proc. Natl. Acad. Sci. U. S. A. 109, 9000–9005. doi: 10.1073/pnas.1118364109.

Deng, W., Ying, H., Helliwell, C. A., Taylor, J. M., Peacock, W. J., and Dennis, E. S. (2011). FLOWERING LOCUS C (FLC) regulates development pathways throughout the life cycle of Arabidopsis. Proc. Natl. Acad. Sci. U. S. A. 108, 6680–6685. doi: 10.1073/pnas.1103175108.

Galvão, V. C., Horrer, D., Küttner, F., and Schmid, M. (2012). Spatial control of flowering by DELLA proteins in Arabidopsis thaliana. Dev. 139, 4072–4082. doi: 10.1242/dev.080879.

Griffiths, J., Murase, K., Rieu, I., Zentella, R., Zhang, Z. L., Powers, S. J., et al. (2006). Genetic characterization and functional analysis of the GID1 gibberellin receptors in *Arabidopsis*. Plant Cell 18, 3399–3414. doi: 10.1105/tpc.106.047415.

Habu, T., Yamane, H., Igarashi, K., Hamada, K., Yano, K., and Tao, R. (2012). 454-pyrosequencing of the transcriptome in leaf and flower buds of Japanese apricot (*Prunus mume* Sieb. et Zucc.) at different dormant stages. J. Japanese Soc. Hortic. Sci. 81, 239–250. doi: 10.2503/jjshs1.81.239.

Hoffmann, M. H. (2005). Evolution of the realized climatic niche in the genus *Arabidopsis* (Brassicaceae). Evolution (N. Y). 59, 1425–1436. doi: 10.1111/j.0014-3820.2005.tb01793.x.

Hyun, Y., Richter, R., Vincent, C., Martinez-Gallegos, R., Porri, A., and Coupland, G. (2016). Multi-layered Regulation of SPL15 and Cooperation with SOC1 Integrate Endogenous Flowering Pathways at the *Arabidopsis* Shoot Meristem. Dev. Cell 37, 254–266. doi: 10.1016/j.devcel.2016.04.001.

Hyun, Y., Vincent, C., Tilmes, V., Bergonzi, S., Kiefer, C., Richter, R., et al. (2019). Plant science: A regulatory circuit conferring varied flowering response to cold in annual and perennial plants. Science (80-.). 363, 409–412. doi: 10.1126/science.aau8197.

Kardailsky, I., Shukla, V. K., Ahn, J. H., Dagenais, N., Christensen, S. K., Nguyen, J. T., et al. (1999). Activation tagging of the floral inducer FT. Science (80-.). 286, 1962–1965. doi: 10.1126/science.286.5446.1962.

Kinoshita, A., and Richter, R. (2021). Genetic and molecular basis of floral induction in Arabidopsis thaliana. J. Exp. Bot. 71, 2490–2504. doi: 10.1093/JXB/ERAA057.

Kobayashi, Y., Kaya, H., Goto, K., Iwabuchi, M., and Araki, T. (1999). A pair of related genes with antagonistic roles in mediating flowering signals. Science (80-.). 286, 1960–1962. doi: 10.1126/science.286.5446.1960.

Kudo, G. (1993). Relationship between flowering time and fruit set of the entomophilous alpine shrub, *Rhododendron aureum* (Ericaceae), inhabiting snow patches. Am. J. Bot. 80, 1300–1304. doi: 10.2307/2445714.

Kudoh, H. (2016). Molecular phenology in plants: In natura systems biology for the comprehensive understanding of seasonal responses under natural environments. New Phytol. 210, 399–412. doi: 10.1111/nph.13733.

Madrid, E., Chandler, J. W., and Coupland, G. (2021). Gene regulatory networks controlled by FLOWERING LOCUS C that confer variation in seasonal flowering and life history. J. Exp. Bot. 72, 4–14. doi: 10.1093/jxb/eraa216.

Mateos, J. L., Madrigal, P., Tsuda, K., Rawat, V., Richter, R., Romera-Branchat, M., et al. (2015). Combinatorial activities of SHORT VEGETATIVE PHASE and FLOWERING LOCUS C define distinct modes of flowering regulation in Arabidopsis. Genome Biol. 16, 1–23. doi: 10.1186/s13059-015-0597-1.

Michaels, S. D., and Amasino, R. M. (1999). *FLOWERING LOCUS C* encodes a novel MADS domain protein that acts as a repressor of flowering. Plant Cell 11, 949–956. doi: 10.1105/tpc.11.5.949.

Munguía-Rosas, M. A., Ollerton, J., Parra-Tabla, V., and De-Nova, J. A. (2011). Meta-analysis of phenotypic selection on flowering phenology suggests that early flowering plants are favoured. Ecol. Lett. 14, 511–521. doi: 10.1111/j.1461-0248.2011.01601.x.

Nagahama, A., Kubota, Y., and Satake, A. (2018). Climate warming shortens flowering duration: a comprehensive assessment of plant phenological responses based on gene expression analyses and mathematical modeling. Ecol. Res. 33, 1059–1068. doi: 10.1007/s11284-018-1625-x.

Pajoro, A., Biewers, S., Dougali, E., Valentim, F. L., Mendes, M. A., Porri, A., et al. (2014). The (r)evolution of gene regulatory networks controlling *Arabidopsis* plant reproduction: A two-decade history. J. Exp. Bot. 65, 4731–4745. doi: 10.1093/jxb/eru233.

Plackett, A. R. G., Powers, S. J., Fernandez-Garcia, N., Urbanova, T., Takebayashi, Y., Seo, M., et al. (2012). Analysis of the developmental roles of the *Arabidopsis* gibberellin 20-oxidases demonstrates that *GA20ox1, −2*, and −*3* are the dominant paralogs. Plant Cell 24, 941–960. doi: 10.1105/tpc.111.095109.

Porri, A., Torti, S., Romera-Branchat, M., and Coupland, G. (2012). Spatially distinct regulatory roles for gibberellins in the promotion of flowering of *Arabidopsis* under long photoperiods. Dev. 139, 2198–2209. doi: 10.1242/dev.077164.

Putterill, J., and Varkonyi-Gasic, E. (2016). FT and florigen long-distance flowering control in plants. Curr. Opin. Plant Biol. 33, 77–82. doi: 10.1016/j.pbi.2016.06.008.

Quiroz, S., Yustis, J. C., Chávez-Hernández, E. C., Martínez, T., Sanchez, M. de la P., Garay-Arroyo, A., et al. (2021). Beyond the genetic pathways, flowering regulation complexity in Arabidopsis thaliana. Int. J. Mol. Sci. 22. doi: 10.3390/ijms22115716.

Rosenzweig, C., Karoly, D., Vicarelli, M., Neofotis, P., Wu, Q., Casassa, G., et al. (2008). Attributing physical and biological impacts to anthropogenic climate change. Nature 453, 353–357. doi: 10.1038/nature06937.

Sasaki, R., Yamane, H., Ooka, T., Jotatsu, H., Kitamura, Y., Akagi, T., et al. (2011). Functional and expressional analyses of *PmDAM* genes associated with endodormancy in Japanese apricot. Plant Physiol. 157, 485–497. doi: 10.1104/pp.111.181982.

Satake, A., Kawagoe, T., Saburi, Y., Chiba, Y., Sakurai, G., and Kudoh, H. (2013). Forecasting flowering phenology under climate warming by modelling the regulatory dynamics of flowering-time genes. Nat. Commun. 4, 1–8. doi: 10.1038/ncomms3303.

Satake, A., Nagahama, A., and Sasaki, E. (2022). A cross-scale approach to unravel the molecular basis of plant phenology in temperate and tropical climates. New Phytol. 233, 2340–2353. doi: 10.1111/nph.17897.

Stuble, K. L., Bennion, L. D., and Kuebbing, S. E. (2021). Plant phenological responses to experimental warming—A synthesis. Glob. Chang. Biol. 27, 4110–4124. doi: 10.1111/gcb.15685.

Sung, S., and Amasino, R. M. (2005). Remembering winter: Toward a molecular understanding of vernalization. Annu. Rev. Plant Biol. 56, 491–508. doi: 10.1146/annurev.arplant.56.032604.144307.

Turck, F., Fornara, F., and Coupland, G. (2008). Regulation and identity of florigen: Flowering Locus T moves center stage. Annu. Rev. Plant Biol. 59, 573–594. doi: 10.1146/annurev.arplant.59.032607.092755.

Vimont, N., Fouché, M., Campoy, J. A., Tong, M., Arkoun, M., Yvin, J. C., et al. (2019). From bud formation to flowering: Transcriptomic state defines the cherry developmental phases of sweet cherry bud dormancy. BMC Genomics 20, 1–23. doi: 10.1186/s12864-019-6348-z.

Vitasse, Y., Ursenbacher, S., Klein, G., Bohnenstengel, T., Chittaro, Y., Delestrade, A., et al. (2021). Phenological and elevational shifts of plants, animals and fungi under climate change in the European Alps. Biol. Rev. 96, 1816–1835. doi: 10.1111/brv.12727.

Whittaker, C., and Dean, C. (2017). The *FLC* locus: A platform for discoveries in epigenetics and adaptation. Annu. Rev. Cell Dev. Biol. 33, 555–575. doi: 10.1146/annurev-cellbio-100616-060546.

Willige, B. C., Ghosh, S., Nill, C., Zourelidou, M., Dohmann, E. M. N., Maier, A., et al. (2007). The DELLA domain of GA INSENSITIVE mediates the interaction with the GA INSENSITIVE DWARF1A gibberellin receptor of Arabidopsis. Plant Cell 19, 1209–1220. doi: 10.1105/tpc.107.051441.

Wilson, R. N., Heckman, J. W., and Somerville, C. R. (1992). Gibberellin is required for flowering in *Arabidopsis thaliana* under short days. Plant Physiol. 100, 403–408. doi: 10.1104/pp.100.1.403.

Wu, G., Park, M. Y., Conway, S. R., Wang, J. W., Weigel, D., and Poethig, R. S. (2009). The Sequential Action of miR156 and miR172 Regulates Developmental Timing in *Arabidopsis*. Cell 138, 750–759. doi: 10.1016/j.cell.2009.06.031.

Yamaguchi, A., Kobayashi, Y., Goto, K., Abe, M., and Araki, T. (2005). *TWIN SISTER OF FT (TSF)* acts as a floral pathway integrator redundantly with FT. Plant Cell Physiol. 46, 1175–1189. doi: 10.1093/pcp/pci151.

Yamane, H., Kashiwa, Y., Ooka, T., Tao, R., and Yonemori, K. (2008). Suppression subtractive hybridization and differential screening reveals endodormancy-associated expression of an *SVP/AGL24-type* MADS-box gene in lateral vegetative buds of japanese apricot. J. Am. Soc. Hortic. Sci. 133, 708–716. doi: 10.21273/jashs.133.5.708.

