## Supplementary material for "The transcriptional changes underlying the flowering phenology shift in response to climate warming": Suppl. Fig. 1

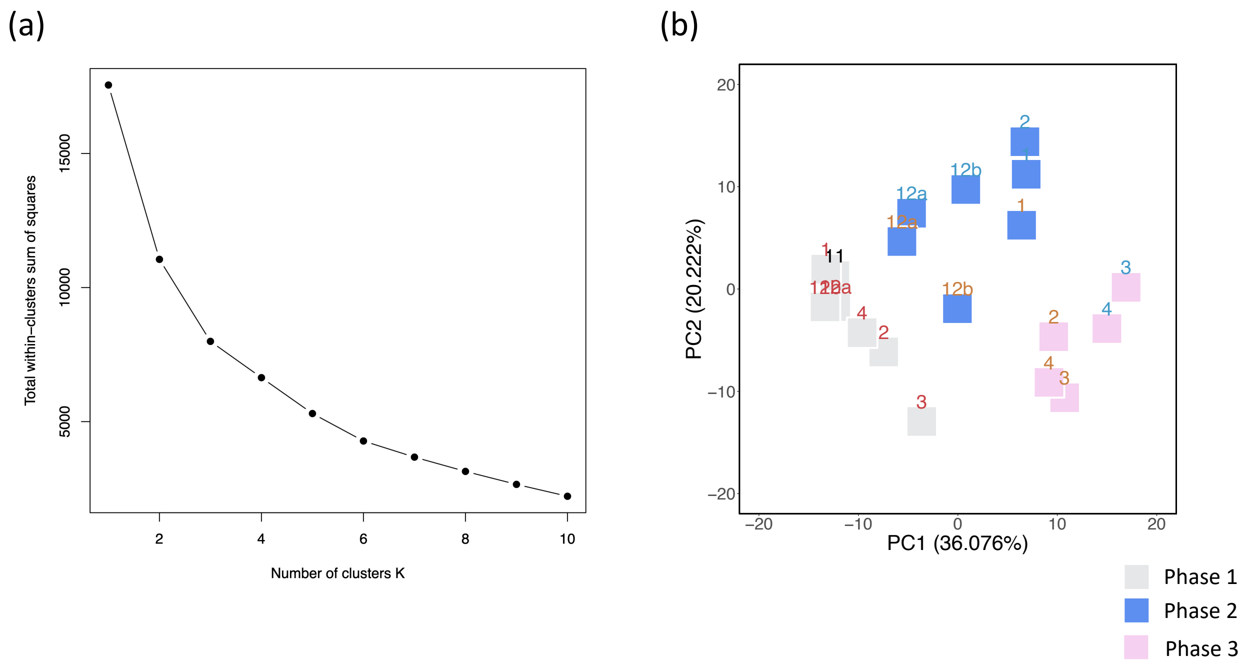


**SUPPLEMENT FIGURE 1 Three phases in the transcriptional profiles based on the non-hierarchical clustering.** (a) Total within-clusters sum of squares by the elbow method. The optimal number of clusters is the point at which the decrease in the total within-clusters sum of squares becomes much smaller as the number of clusters increases. In this case, the optimal number of clusters is 3. **(b)** A plot of PC2 versus PC1 resulted from the PCA of the 293 genes. Plot colors indicate the gene expression profiles for phase 1(gray), phase 2 (blue), and phase 3 (pink). The division of the three phases is based on the result of non-hierarchical clustering. Letter colors indicate sampling sites: the lab (black), the northern site (blue), the southern site (orange), and the subtropical site (red). The numbers in brackets represent the explained variance.

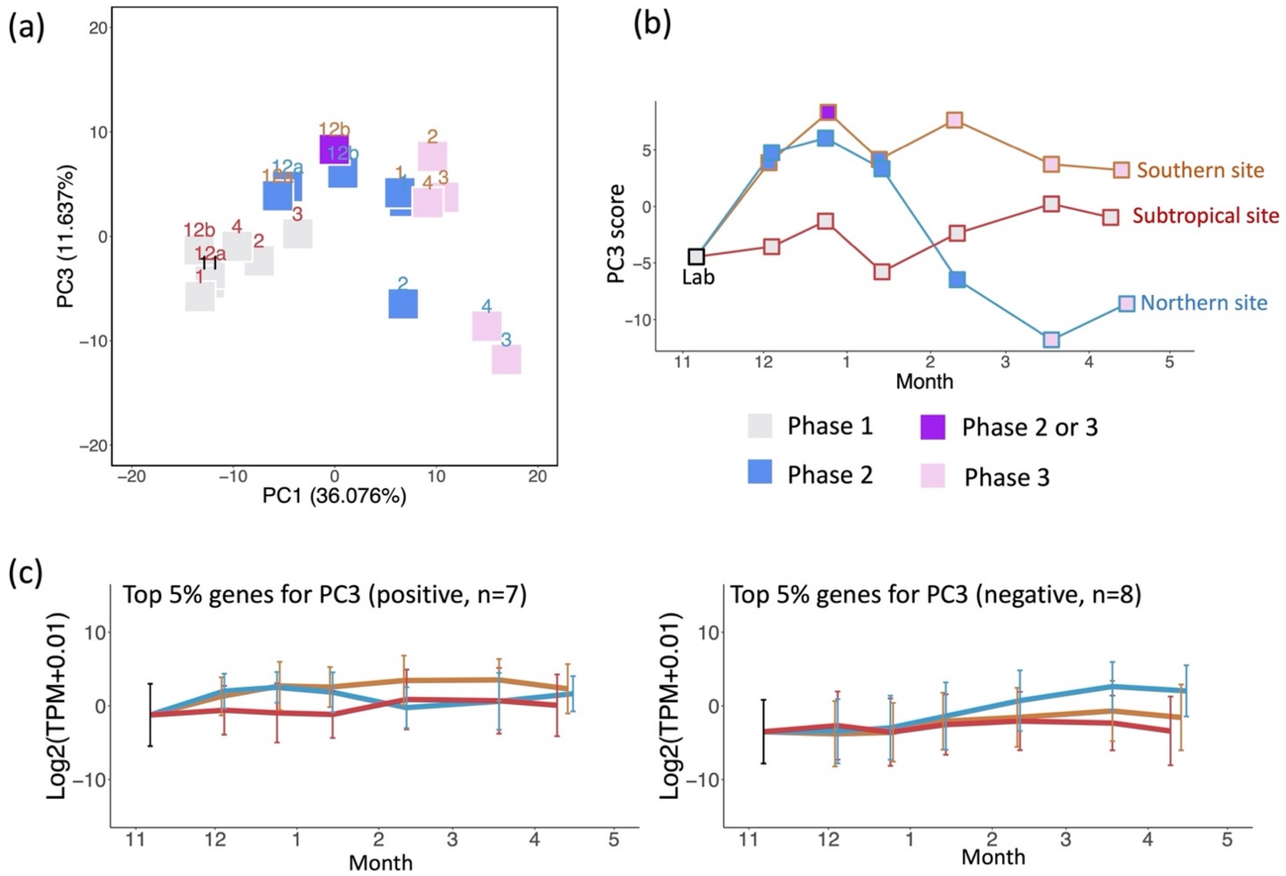


**SUPPLEMENT FIGURE 2 Minor axis of multivariate molecular phenology.** (a) A plot of PC3 versus PC1 resulted from the PCA of the 293 genes. Letter colors indicate sampling sites: the northern site (blue), the southern site (orange), and the subtropical site (red). Plot colors indicate the gene expression profiles for phase 1(gray), phase 2 (blue), phase 2 or 3 (purple), and phase 3 (pink). The numbers in brackets represent the explained variance. (b) A Plot of the PC3 scores against the month. (c) The expression profile of the genes, specifically of those genes with either positive or negative eigenvectors in the top 5% of the largest absolute values of eigenvectors in PC3, respectively. The average (line) ± s.d. (error bar) of those genes is shown.

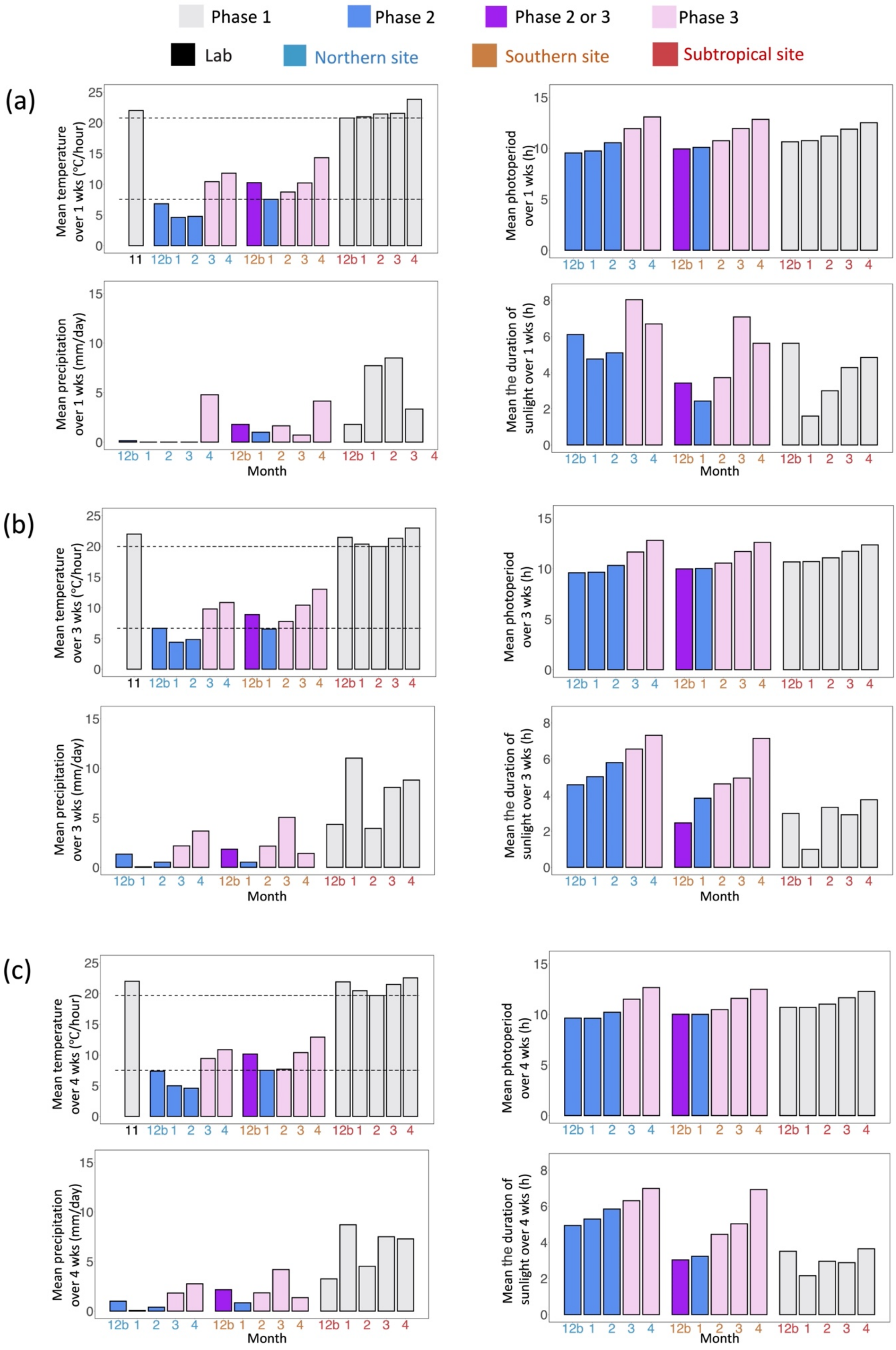


**SUPPLEMENT FIGURE 3 Molecular phenology of flowering-time genes was associated with seasonal changes in temperature.**

The mean temperature, photoperiod, precipitation, and duration of sunlight over a period of one (a), three (b), and four (c) weeks preceding the sampling dates are illustrated. The Dashed lines indicate 20.8°C, 20.0°C, and 19.7°C above which temperature the phase 1 profiles in gene expression emerge, and 7.6°C, 6.7°C and 7.5°C below which temperature the phase 2 profiles in gene expression emerge in (a), (b), and (c), respectively. Note that the plants were not directly impacted by rainfall as they were shielded by a roof at the subtropical site.

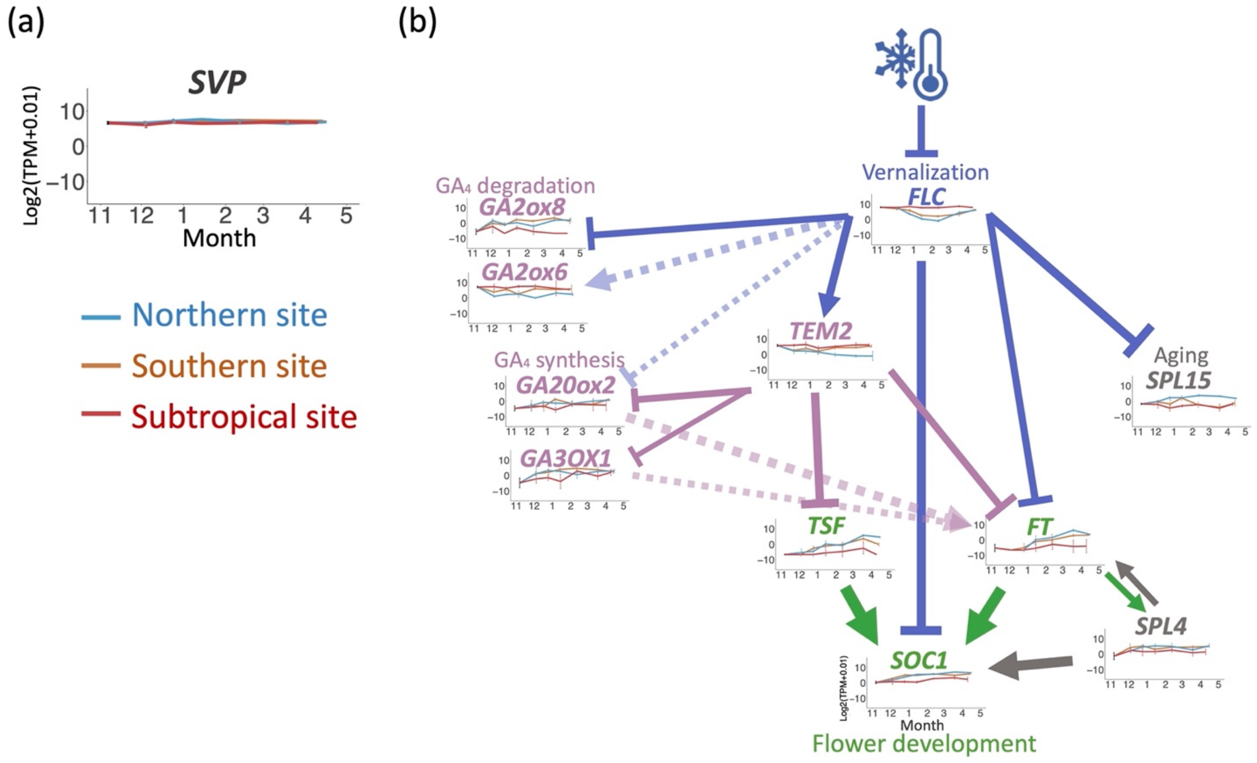


**SUPPLEMENT FIGURE 4 Seasonal transcriptional changes in the network of flowering-time genes.** (a) The expression profile of *AhgSVP* (*SHORT VEGETATIVE PHASE*). The average (line) ± s.d. (error bar) is shown (n = 3; biological replicates). (b) Flowering gene network using the top 5 % of genes with the largest absolute values of eigenvectors for PC1. The color depends on which pathway the gene belongs to blue for the vernalization, pink for the hormonal pathway, grey for the aging pathway, and green for the flower development pathway. The dotted lines with relatively lighter colors indicate indirect effects based on the findings of *A. thaliana* (Mateos et al., 2015; Bouché et al., 2016). Arrow and T-bar indicate positive and negative regulation based on the slope of linear regression analysis.
